# Specific telomere protection ensured by FOXO3a upon genotoxic stress and during aging

**DOI:** 10.1101/2021.08.04.454762

**Authors:** Maria Sol Jacome Burbano, Jérome Robin, Serge Bauwens, Marjorie Martin, Emma Donati, Lucia Martínez, Sabrina Sacconi, Frédérique Magdinier, Eric Gilson

## Abstract

Longevity is determined by diverse signaling pathways including telomere protection and homeostasis master regulators like FOXO3a. We previously showed that the telomeric repeat binding factor 2 (TRF2) expression decreases with age in human skeletal muscle and that, surprisingly, its loss in myofibers does not trigger telomere deprotection. We reveal here that in *TERF2*-compromised myotubes, FOXO3a is recruited to telomeres where it acts as a protective factor against ATM-dependent DNA damage activation. Moreover, we show that FOXO3a-telomere association increases with age in human skeletal muscle biopsies. In mitotic fibroblasts, the telomere protective properties of FOXO3a are operative if the cells are treated with bleomycin. The telomere function of FOXO3a does not require its Forkhead DNA binding domain but the CR2C. Overall, these findings demonstrate a direct connection between two key longevity pathways, FOXO3a and telomere protection. This unveils an unexpected higher level of integration in the regulation of longevity signaling pathway.

## Introduction

Aging can be defined as a progressive failure of homeostasis and organ function that increases susceptibility to several diseases and limits longevity (Singh *et al*, 2019). An extraordinary diversity of aging trajectories exists in nature (Jones *et al*, 2014), which can be explained by the effects of natural selection aimed at maximizing fitness in a given environmental context (Partridge & Barton, 1993). Mechanistically, aging is caused by limitation of somatic maintenance and stress response mechanisms, leading to a gradual increase in molecular and cellular damage along with senescent cell accumulation (Gorgoulis *et al*, 2019; Schumacher *et al*, 2021). The rate of deterioration is determined by interconnected biological processes, known as aging hallmarks, which are regulated by environmental and genetic factors (López-Otín *et al*, 2013). Pro-longevity genes, protecting against aging hallmarks, have been identified in human populations through genome-wide association studies. For example, several studies have consistently identified *FOXO3a* as a pro-longevity master transcription factor that serves as a homeostasis regulator (Webb & Brunet, 2014; Soerensen *et al*, 2015). Meanwhile, the possible existence of one or several developmentally regulated clocks that orchestrate the aging hallmarks remains unclear. In humans and other vertebrates, telomeres may represent such an aging clock, as their DNA length is programmed to shorten in somatic cells during embryonic development (Wright *et al*, 1996; Gilson & Géli, 2007) and upon stress exposure, accounting in part for the disparate trajectories of aging (Epel *et al*, 2004; Jacome Burbano & Gilson, 2021). Moreover, dysregulation of telomeres is associated with rare progeroid syndromes (Armanios & Blackburn, 2012) and a broad spectrum of age-related pathologies in the general population (Martínez *et al*, 2017), while their reinforcement extends the lifespan in mouse models (Muñoz-Lorente *et al*, 2019). This central position of telomeres as an aging regulator might stem from their interconnection to several aging hallmarks such as genome stability (de Lange, 2018), senescence (Abdallah *et al*, 2009), oxidative stress (Barnes *et al*, 2019), and mitochondrial integrity (Sahin *et al*, 2011; Robin *et al*, 2020).

Many telomere functions are performed by a capping protein complex called shelterin. It consists of proteins directly bound to telomeric DNA (in humans, TRF1, TRF2, and POT1) and other proteins that create a bridge between the duplex DNA and the 3’ overhang (in humans, TPP1/ACD and TIN2) or are simply bound to other subunits (in humans, RAP1 bound to TRF2). Among shelterin subunits, TRF2 may mediate important regulatory functions during aging, as it interconnects various aging hallmarks (genome stability, senescence, mitochondria, and immunity), is downregulated during aging in multiple tissues (Robin *et al*, 2020; Wagner *et al*, 2017), and plays pivotal roles in aging in model organisms (Kishi *et al*, 2008; Alder *et al*, 2015; Morgan *et al*, 2019). The multiple roles of TRF2 rely on its capacity to blunt the DNA damage response (DDR) ataxia–telangiectasia mutated (ATM) checkpoint and non-homologous DNA repair at chromosomal termini, in addition to genomewide transcriptional functions (Ye *et al*, 2014). For instance, TRF2 regulates the expression of glycocalyx genes involved in immunosurveillance (Cherfils-Vicini *et al*, 2019), the *Sirt3* sirtuin gene required for mitochondrial integrity (*20*) and the *hTERT* telomerase gene (Kim *et al*, 2016; Sharma *et al*, 2021).

Strikingly, in skeletal muscle fibers, the absence of TRF2 does not lead to detectable telomere damage, suggesting that an alternative telomere protective mechanism exists in these cells (Robin *et al*, 2020). Here, we demonstrate that FOXO3a binds to and protects telomeres of post-mitotic skeletal muscle fibers upon *TERF2* downregulation. The role of FOXO3a in telomere protection is conserved in dividing cells upon genotoxic stress. These results reveal a connection between two important longevity pathways.

## Results

### 1. FOXO3a is required for telomere protection upon *TERF2* downregulation in myotubes

In accordance with our previous findings (Robin *et al*, 2020), *TERF2* knockdown (KD) in human myotubes did not trigger DDR specifically at telomeres, as measured by the ratio of 53BP1 nuclear foci associated with telomeres (designated specific telomere dysfunction-induced foci or sTIF), which led to a high cellular level of reactive oxygen species (ROS) (Fig. 1A–C; Fig. EV1 A, B). Moreover, *TERF2* downregulation in myotubes activates FOXO3a (Robin *et al*, 2020), a longevity factor involved in the oxidative stress response and muscle homeostasis (Sandri *et al*, 2004; Mammucari *et al*, 2007; Zhao *et al*, 2007; Milan *et al*, 2015). Therefore, we investigated whether FOXO3a could act at telomeres in *TERF2-com*promised myotubes. Interestingly, downregulation of *FOXO3a* expression in myotubes led to a specific loss of telomere protection against DDR only with simultaneous *TERF2* downregulation (Fig. 1A–C). Further, treatment with chemical inhibitors of the DDR kinases ATM (KU-55933) or ATR (ATR; VE-821) revealed that the telomere-specific damage triggered by double downregulation of *TERF2* and *FOXO3a* in myotubes was ATM dependent (Figure 1D).

**Fig. 1:**
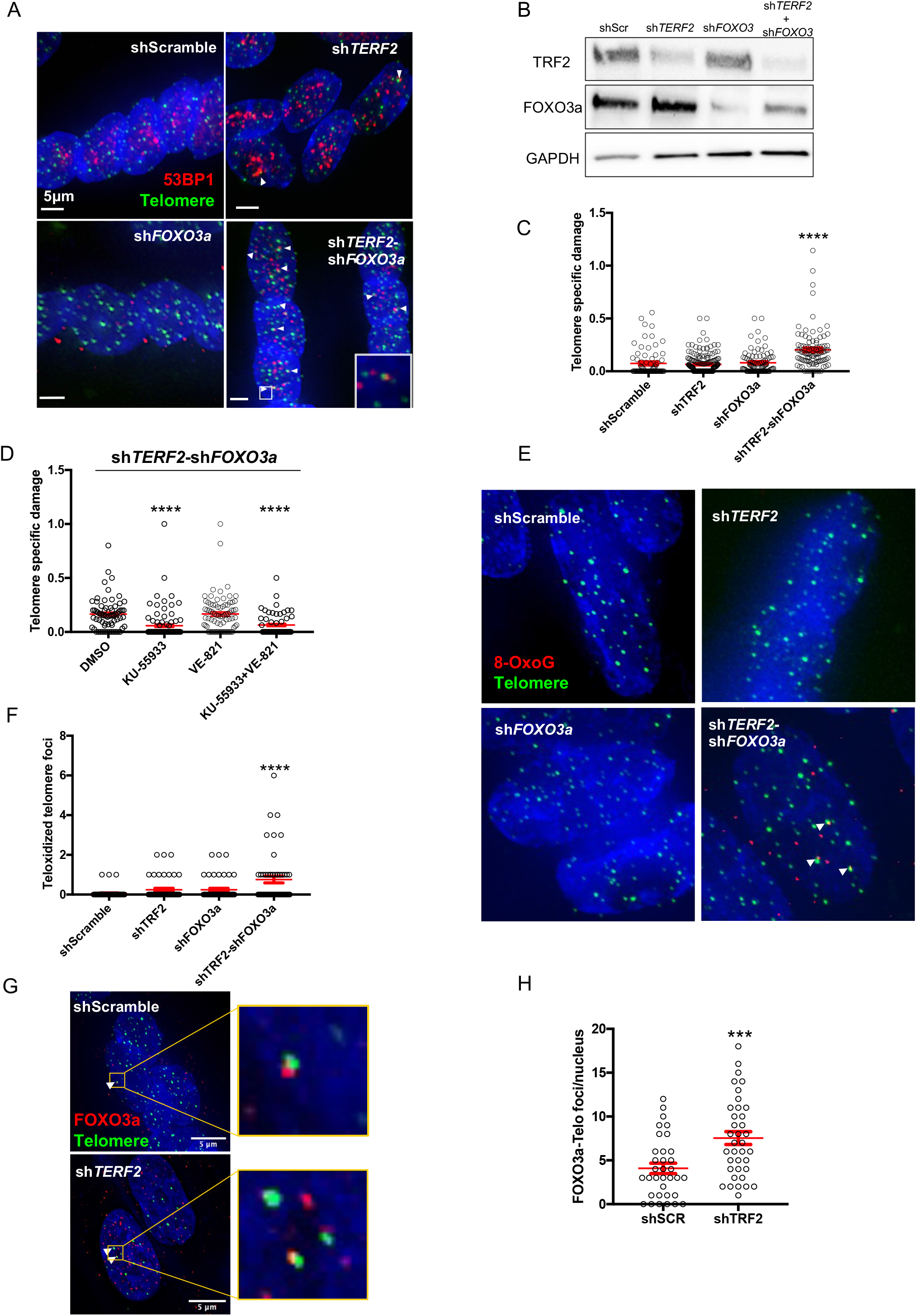
FOXO3a directly protects telomeres upon TRF2 downregulation in myotubes. **(A)** Telomere Induced damage Foci assay in transduced myotubes (10 days), using a telomeric PNA probe (green) and 53BP1 (red) staining. Colocalizations pointed by an arrow, indicate DDR activation at telomeres. 30-40 nuclei were analyzed per replicate and per condition, n=4. **(B)** Western Blotting showing *TERF2* and *FOXO3a* downregulation. In *shTERF2* myotubes, FOXO3a expression is increased. **(C)** Ratio between the number of TIFs and the total number of 53BP1 within each nucleus, corresponding to the specific telomere damage (sTIF). shScramble vs *shTERF2-shFOXO3a*, *p-value* <0.000001. **(D)** Telomeric specific damage (TIFs/53BP1 ratio) in myotubes inhibited for ATM and/ or ATR. 5 days after differentiation and 5 days before harvesting, ATM or ATR was inhibited using either KU-55933 (10mM) or VE-821 (10mM), respectively. Fresh media with chemical inhibitors was added every 2 days. Bars represent SEM of two biological replicates. *p-value* <0.000001. **(E)** Immunofluorescence detection of 8-Oxoguanosine (red), combined with telomeric FISH probe (green) in transduced myotubes and quantified in **(F)**. Oxidated telomeres (TOFs) were analyzed by colocalization of the green and red signals. Bars represent SEM of two biological replicates (approximately 35 nuclei). shScramble vs *shTERF2-shFOXO3a, p-value* <0.000001. **(G)** Immunofluorescence detection of FOXO3a (red), combined with telomeric FISH probe (green) in myotubes transduced with sh*TERF2*. Colocalization are shown by a white arrow and a zoom is shown at the left. Quantification of FOXO3-Telomere foci per nucleus in H). *p-value*= 0,0008. Statistical analyses were performed using Kruskal-Wallis’ multiple comparisons test (\**p* < 0.05, \*\**p* < 0.001, \*\*\**p* < 0.001, \*\*\*\**p* < 0.0001).

In addition, downregulation of *FOXO3a* expression did not increase the high ROS level induced by inhibition of *TERF2* expression (Fig. EV1 C–F). Similarly, the expression of a panel of oxidative stress-related genes (*NRF2*, *GSS*, *BNIP3*, *PRDX1*, *SOD2*, and *CAT*) was not impaired by *FOXO3a* downregulation in *TERF2*-compromised myotubes (Fig. EV1 G–L). Therefore, telomere deprotection triggered by *FOXO3a* expression inhibition in *TERF2*-compromised myotubes cannot be explained simply by a global increase in ROS or a decrease in oxidative defense.

Next, because telomeres are G-rich and particularly sensitive to oxidation through the formation of 8-oxoguanosine (8-oxoG), we examined whether FOXO3a plays a role in this specific oxidation process. To that end, we quantified colocalization events between telomeric DNA foci marked with a fluorescent PNA probe and 8-oxoG foci revealed with specific antibodies (known as teloxidized telomere foci or TOF). Echoing sTIF, only double downregulation of *TERF2* and *FOXO3* increased the mean number of TOFs per nucleus (Fig. 1E, F).

Our results suggest that FOXO3a is capable of decreasing oxidation and ATM-dependent DDR activation specifically at telomeres in a post-mitotic context (myotubes). As this telomeric role of FOXO3a is revealed in the context of oxidative stress triggered by *TERF2* downregulation, we hypothesized that FOXO3a provides an alternative strategy for telomere protection in the presence of genotoxic stress.

### 2. FOXO3a localizes to myotube telomeres

Next, we investigated whether the telomere protective role of FOXO3a is directed through assessment of its putative association with telomeres. On average, 17 FOXO3a foci were detected in untreated myotube nuclei, of which four specific colocalizations with telomeres were observed using a PNA probe (Fig. 1G, H). Upon *TERF2* downregulation, in parallel with nuclear translocation of activated FOXO3a (Fig. EV1 M), we measured a roughly two-fold increase in such colocalization events (Fig. 1G, H). To validate this interaction between telomeres and FOXO3a, we performed a proximity ligation assay (PLA) using TRF1 and FOXO3a as potential partners. Moreover, we assessed FOXO3a enrichment at telomeres through chromatin immunoprecipitation (ChIP) with slot-blotting quantification using a radiolabeled telomeric DNA probe (Fig EV1 N–Q). We concluded that FOXO3a plays a direct role in telomere protection in myotubes upon *TERF2* downregulation.

Based on these results, we investigated the interaction in an *in vivo* context using our previously published set of muscle biopsies (Robin *et al*, 2020). We confirmed the presence of FOXO3a at telomeres using co-immunoprecipitation (co-IP) with an antibody against TRF2 based on FOXO3a immunoblotting (Fig. EV1 R). Notably, the level of co-IP increased with age (Fig. EV1 S, T). This result suggests an important role of the FOXO3a–telomere association during skeletal muscle aging. Older muscle biopsies contain lower levels of TRF2 (Robin *et al*, 2020), and therefore the increase in TRF2–FOXO3a co-IP with age can be interpreted as a high level of oxidative stress triggering the association of FOXO3a with the residual TRF2 present at telomeres, in accordance with observations in TERF2-compromised myotubes.

### 3. FOXO3a acts synergistically with TERF2 to protect telomeres in transformed fibroblasts

Next, we investigated whether the telomeric protection function of FOXO3a is conserved in another cellular context (*i.e*., mitotic cells). To this end, we used BJ-HELT transformed fibroblasts derived from human foreskin cells expressing *hTERT* and *SV40 early region* (large-T and small-t antigens) genes. In these cells, downregulation of *FOXO3a* did not lead to specific telomere damage, whereas *TERF2* downregulation caused such damage, as expected (Fig. 2A, B). Notably, double downregulation of *FOXO3a* and *TERF2* showed synergistic effects, exacerbating telomere damage (Fig. 2A, B). This situation is reminiscent of myotubes, wherein the telomere protective function of FOXO3a is revealed only upon *TERF2* downregulation.

**Fig. 2:**
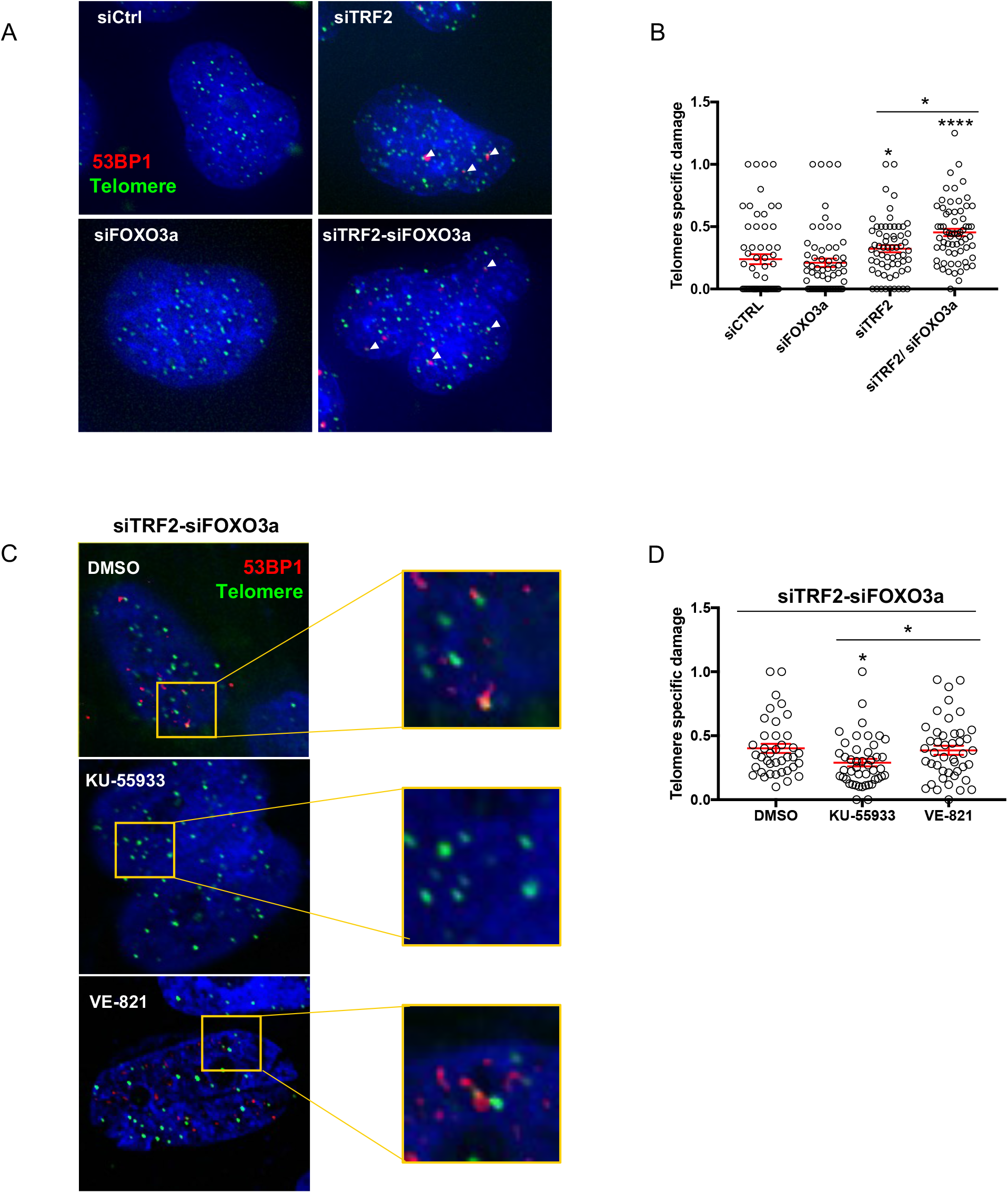
FOXO3a protects telomeres upon *TERF2* KD in BJ-HELT fibroblasts. **(A)** Telomere Induced damage Foci assay in transfected BJ-HELT (72h siRNA), using a telomeric PNA probe (green) and 53BP1 (red) staining. 53BP1 - Telomeres colocalizations are pointed by a white arrow. **(B)** Quantification of sTIF observed in **(A)**. 30-40 nuclei were analyzed in three biological replicates. siCtrl *vs* siTRF2 *p-value=* 0,008304 and siCtrl *vs* siTRF2-siFOXO3a *p-value* <0.000001. **(C)** TIF assay in siTRF2-siFOXO3a transfected BJ-HELT cells (72h siRNA, PNA-telomere probe in green and 53BP1 in red), inhibited for ATM or ATR. Inhibitors were added 24h before cell harvest: KU-55933 (10mM) or VE-821 (10mM), respectively. Zoom at the left. **(D)** sTIF quantification. Bars represent SEM of two biological replicates. DMSO *vs* KU-55933 *p-value* =0,0112. Statistical analysis were performed using Kruskal-Wallis’ multiple comparisons test; α=0.05)

Therefore, we explored whether FOXO3a-dependent telomere protection is specific to *TERF2-*compromised cells by repeating the previous experiment with sequential downregulation of each shelterin subunit (Fig. EV2 A–D). As expected, individual downregulation of *TERF1*, *TERF2*, *TPP1*, *POT1*, or *TINF2* induced specific telomere damage. When coupled downregulation with *FOXO3a* was tested via RNA interference, only the *TERF2*–*FOXO3a* KD condition increased the level of specific telomere damage. Thus, FOXO3a appears to be an alternative protective factor against telomere damage induced by TRF2 loss. This finding is in accordance with the ATM-dependent damage observed with concomitant downregulation of *TERF2* and *FOXO3a* in myotubes (Fig. 1D) and BJ-HELT cells (Fig. 2C, D). In contrast to myotubes, we failed to detect any difference in FOXO3a expression or nuclear translocation upon *TERF2* downregulation (Fig. EV2 E–I) in BJ-HELT cells, indicating that FOXO3a-mediated telomere protection in *TERF2*-compromised BJ-HELT fibroblasts does not result from activation due to oxidative stress.

### 4. FOXO3a specifically protects telomeres from ATM-recognized damage

To determine the type of telomere damage that FOXO3a specifically protects against, we exposed BJ-HELT cells to a set of genotoxic stressors known to cause DNA damage that may be repaired by various processes (H2O2, bleomycin, paraquat, sodium arsenite, and ultraviolet radiation) (Fig. 3A; Fig. EV3 A, B). Only bleomycin-treated cells showed an effect of *FOXO3a* downregulation on telomere protection (Fig. 3A, B). With *TERF2* downregulation, FOXO3a specifically protected against ATM-recognized damage in bleomycin-treated BJ-HELT cells (Fig. 3C). These results are reminiscent of previous work reporting that FOXO3a protects the genome against instability caused by double-strand breaks (White *et al*, 2020). We conclude that FOXO3a protects against multiple types of ATM-recognized damage at telomeres.

**Fig. 3:**
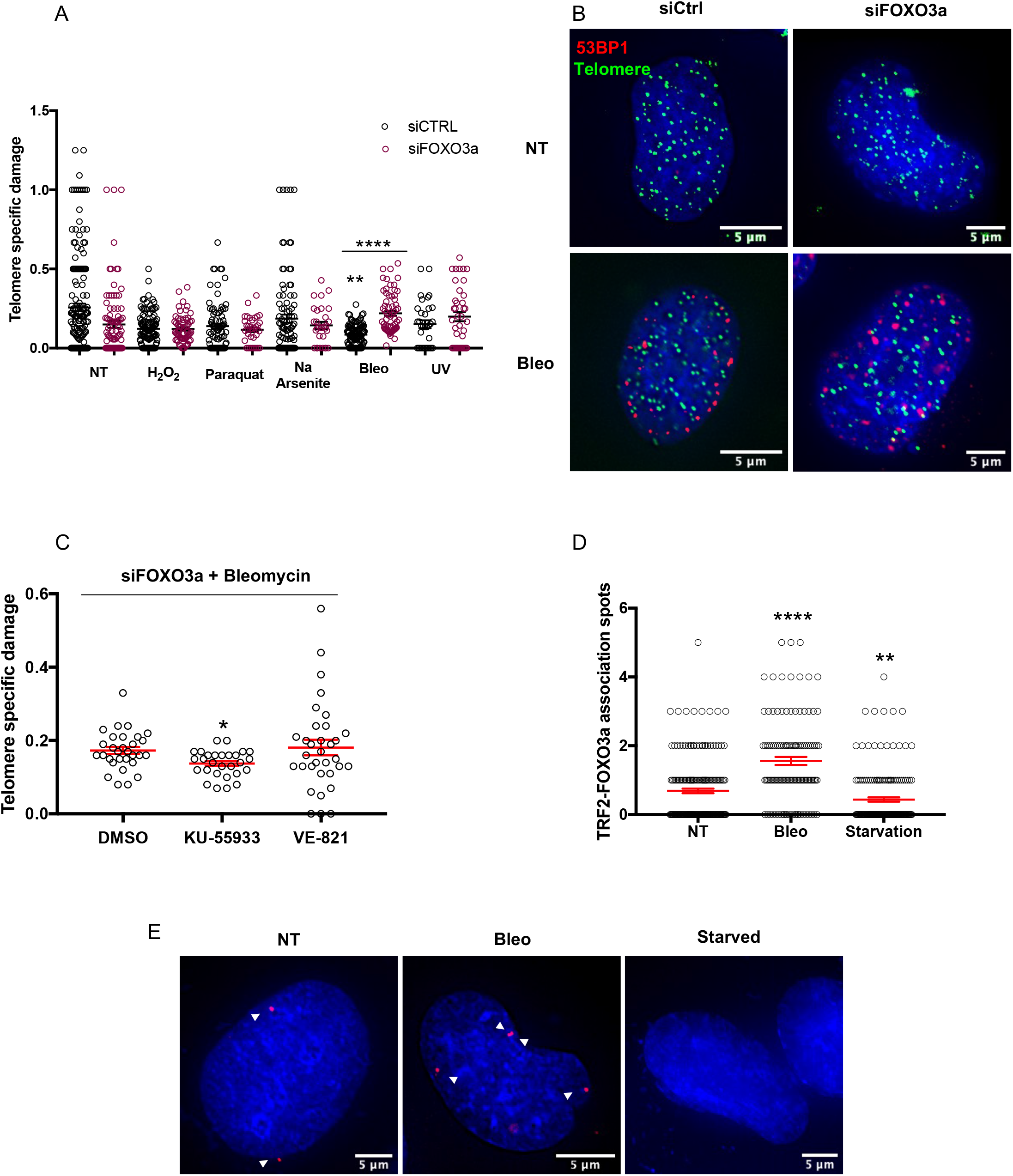
FOXO3a specifically protects telomeres from ATM-recognized damages. **(A)** Telomere specific damage (sTIF) in transfected BJ-HELT cells with siControl (black circles) or siFOXO3a (pink circles) and exposed to different stressors. Statistical analyses were performed using Kruskal-Wallis’ multiple comparisons test (\**p* < 0.05, \*\**p* < 0.001, \*\*\**p* < 0.001, \*\*\*\**p* < 0.0001) in biological triplicates. **(B)** Representative images of TIFs in non-treated or bleomycin exposed BJ-HELT cells downregulated or not for FOXO3a. Immunofluorescence of 53BP1 (red) and telomeric PNA-FISH probe (green). **(C)** sTIF in siFOXO3a-BJ-HELT cells, treated with bleomycin (50 μg/mL, 24hr) and inhibited for ATM (KU-55933) or ATR (VE-821). Two biological replicates analyzed with a Kruskal-Wallis’ multiple comparisons test (DMSO vs KU-55933, *p-value*= 0,0464). **(D)** PLA showing TRF2-FOXO3a association in non-treated BJ-HELT or exposed to bleomycin (50 μg/mL, 24hr) or under caloric restriction. NT *vs* Bleo (*p-value* <0.000001) and NT *vs* Starvation (*p-value*=0,007385) were analyzed using a Kruskal-Wallis’ multiple comparisons in biological triplicates. **(E)** Representative images of TRF2-FOXO3a PLA in E). PLA red foci are pointed by white arrows.

Next, we asked whether, as in myotubes, FOXO3a plays a direct telomere protective role in BJ-HELT cells. We detected TRF2–FOXO3a associations in bleomycin-treated cells using a PLA (Fig. 3D, E; Fig. EV3 C, D). Such associations cannot be attributed only to the partial nuclear translocation of FOXO3a observed upon bleomycin treatment (Fig. EV3 E, F) because starvation, which causes massive localization of FOXO3a in the nucleus, did not increase PLA signals (Fig. 3D, E).

### 5. The CR2C domain of FOXO3a is essential for its role in telomere protection

Finally, we investigated which domain of the FOXO3a protein is involved in telomere protection. To this end, we constructed a battery of lentiviral vectors expressing various truncated forms of FOXO3a: ΔCR1, ΔFH, ΔCR2C, and ΔCR3 (Fig. 4A). Through downregulation of *FOXO3a* via RNA interference in cells overexpressing either wild-type (WT) or truncated FOXO3a, we obtained cells expressing nearly physiological levels of FOXO3a or its truncated forms (Fig. 4B; Fig. EV4 A, B). Importantly, expression of WT FOXO3a was sufficient to restore telomere protection in bleomycin-treated cells, indicating that the small interfering RNA pool we used to downregulate *FOXO3a* expression did not cause off-target effects related to telomere protection (Fig. 4C). Notably, expression of the truncated form with deletion of the Forkhead (FH) DNA-binding domain also rescued telomere protection. Among other truncated forms, only the form with deletion of the CR2C domain was unable to rescue telomere protection (Fig. 4C; Fig. EV4 C, D), and this form also reduced FOXO3a telomeric localization (Fig. 4D). We conclude that FOXO3a exhibits telomere protective properties uncoupled from its canonical role as a DNA-binding transcription factor, but that is dependent on the CR2C domain.

**Fig. 4:**
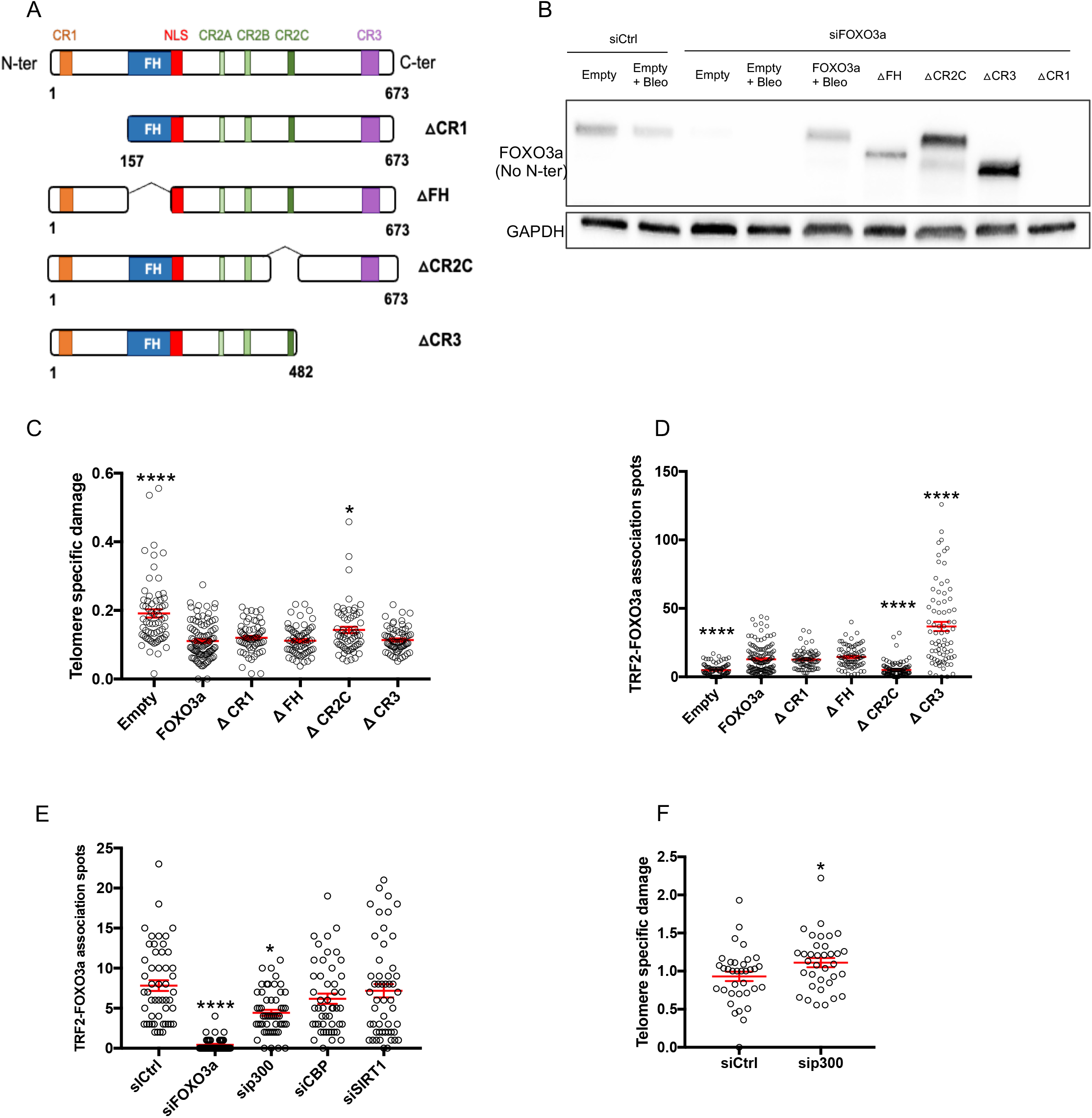
FOXO3a CR2C domain is required for telomere protection. **(A)** FOXO3a full-length structure (top) showing relevant domains and truncated mutant forms (ΔCR1, ΔFH, ΔCR2C and ΔCR3). FOXO3a is composed by three conserved regions: CR1, CR2 and CR3 known to interact with several proteins. CR2 is divided in three subregions: CR2A, CR2B and CR2C. FOXO3a contains a Forkhead (FH) or DNA binding domain, that recognizes a TGTTTAC consensus DNA sequence, followed by a nuclear localization sequence (NLS). **(B)** Western blotting showing endogen FOXO3a downregulation, FOXO3a fulllength rescue and expression of FOXO3a truncated proteins. FOXO3a-Cell Signaling Technologies #2497 monoclonal antibody does not recognize ΔCR1 form because its epitope surrounds Glu50 (N-terminal). Cells were transduced 5 days with lentiviruses and the last 3 days transfected with siRNA. GAPDH was used as loading control. **(C)** sTIF quantification in siFOXO3a + truncated FOXO3a overexpression in BJ-HELT treated with bleomycin (50 μg/mL, 24hr), n=3. Conditions were compared to siFOXO3a-FOXO3a+Bleo using a Kruskal-Wallis’ multiple comparisons test. siFOXO3a-FOXO3a+Bleo *vs* siFOXO3a-△CR2C+Bleo*, p-value*=0,0345. **(D)** PLA showing TRF2-FOXO3a association in siFOXO3a + truncated FOXO3a overexpression in BJ-HELT treated with bleomycin (50 μg/mL, 24hr), n=3. **(E)** TRF2-FOXO3a PLA in transfected BJ-HELT cells treated with bleomycin, n=2. Statistical analyses were performed using Kruskal-Wallis’ multiple comparisons test (\**p* < 0.05, \*\**p* < 0.001, \*\*\**p* < 0.001, \*\*\*\**p* < 0.0001). **(F)** Telomere specific damage upon p300 inhibition. siCtrl and sip300 transfected BJ-HELT cells were treated with bleomycin. Means ± SEM of two replicated are shown, Man-Whitney U test (*p-value=* 0,0346).

The CR2C domain interacts with CBP/p300 and SIRT1 proteins. Interestingly, p300 has been shown to bind and acetylate TRF2 and to be essential for telomere protection in HT1080 cells (Her & Chung, 2013), whereas SIRT1 binds to telomeres and regulates the maintenance of telomere length (Palacios *et al*, 2010; De Bonis *et al*, 2014). Therefore, p300 and SIRT1 are promising candidates that may contribute to the telomere protective properties of FOXO3a. Thus, we downregulated the expression of *p300*, *CBP*, and *SIRT1* in bleomycin-treated cells. Only downregulation of *p300* weakened the TRF2–FOXO3a association (Fig. 4E). Moreover, its downregulation increased telomere-specific damage in bleomycin-treated cells, mimicking the effects of *FOXO3a* downregulation (Fig. 4F; Fig. EV4 E, F). Whereas the TRF2 level was previously reported to decrease upon p300 inhibition (Her & Chung, 2013), we observed no difference in TRF2 protein levels between control and p300-compromised cells (Fig. EV4 G, H).

Overall, our results demonstrate a direct role of FOXO3a in protecting telomeres against ATM-recognized damage through a complex mediated by p300.

## Discussion

Mammalian FOXO transcription factors, homologs of DAF-16, which regulates longevity in *Caenorhabditis elegans*, play a key role in regulating cell homeostasis through upregulation of genes associated with oxidative stress resistance, metabolism, apoptosis, cell cycle arrest, aging, and autophagy (Webb & Brunet, 2014). Here, we reveal a novel function of FOXO3a in providing specific protection to telomeres against a genotoxic insult. Downregulation of *FOXO3a* in unchallenged cells has little or no effect on telomere protection, while FOXO3a uncaps telomeres in cells experiencing genotoxic stress. This conditional protective role of FOXO3a was observed in post-mitotic muscle fibers upon *TERF2* downregulation, leading to mitochondrial dysfunction and telomere oxidation, as well as in transformed fibroblasts upon *TERF2* downregulation combined with treatment with the clastogenic agent bleomycin.

The protective function of FOXO3a on telomeres cannot be explained simply by its general roles in DNA repair and oxidative stress (White *et al*, 2020; Tran *et al*, 2002; Brunet *et al*, 2004), but instead indicates specific properties affecting only telomeres. First, through normalization of the rate of telomere damage to the total amount of nuclear damage, telomeres are better protected than the rest of the genome upon *FOXO3a* downregulation. Second, in *TERF2*-compromised myotubes, the rate of oxidized telomeres increased markedly upon *FOXO3a* downregulation, while the total amount of ROS remained unchanged. Third, a subset of FOXO3a proteins are associated with telomeres, as revealed through various techniques, including colocalization with telomeric PNA probe foci, PLA foci between FOXO3a and either TRF1 or TRF2, ChIP revealed by telomere slot blotting, and co-IP with TRF2. In myotubes, the telomere–FOXO3a association increased upon *TERF2* downregulation, in parallel with an increase in the number of FOXO3a nuclear foci. In transformed fibroblasts, this association also increased upon bleomycin treatment. Notably, this telomere association may be uncoupled from FOXO3a nuclear translocation. Indeed, starvation, which causes massive translocation of FOXO3a into the nucleus, was not correlated with an increase in telomere association; conversely, bleomycin treatment led to increased recruitment of FOXO3a at telomeres in the absence of detectable nuclear translocation. These findings suggest that FOXO3a must undergo specific post-translational modification in response to genotoxic stress to associate with telomeres.

The telomere protective role of FOXO3a is conserved upon deletion of its FH DNA-binding domain or its C-terminal CR3 transactivation domain. These results rule out the possibility of FOXO3a acting at telomeres through its canonical role as a DNA-binding transcription factor. Moreover, deletion of CR3 increases the capacity of FOXO3a to associate with telomeres. This finding suggests competition between the major transactivation domain of FOXO3a and its telomere-specific functions. Deletion of CR3 may also cause FOXO3a to exhibit an open conformation (Wang *et al*, 2008) that is more prone to telomere association. The CR3 domain was reported to bind and activate ATM (Tsai *et al*, 2008; Adamowicz *et al*, 2016), making a protective role of FOXO3a at telomeres via its interaction with ATM unlikely. In contrast to the FH and CR3 domains, deletion of CR2C impairs the telomere protective role of FOXO3a. CR2C exhibits transactivation activities and mediates the associations of FOXO3a with SIRT1 (Nakae *et al*, 2006) and with CBP/p300 (Senf *et al*, 2011; Wang *et al*, 2012). Among *SIRT1*, *CBP*, and *p300*, only downregulation of *p300* impaired the TRF2–FOXO3a association, as revealed by PLA. In accordance with involvement of a FOXO3a–p300 complex in telomere protection, p300, but not CBP, was associated with telomeres (Cubiles *et al*, 2018), raising the possibility that acetylation mediated by p300 may promote the repair of damaged telomeres. This model requires substantiation, particularly through determining the targets of FOXO3a–p300 on telomeres. One candidate is TRF2, which contains a p300 binding site (Her & Chung, 2013). These findings raise the question of the type of damage that causes FOXO3a to be efficiently signaled for telomere repair. Our results suggest that such damage is recognized through the ATM-dependent DDR pathway.

The findings of this study establish a direct link between two key longevity hallmarks, FOXO3a (Martins *et al*, 2016) and telomere protection (Foley *et al*, 2018; Wirthlin *et al*, 2018; Augereau *et al*, 2021). This connection is likely involved in muscle aging. Indeed, telomere protection relies on FOXO3a in *TERF2* depleted muscle fibers, a situation occurring during muscle aging (Robin *et al*, 2020). Furthermore, FOXO3a– telomere increased association with age based on human biopsies. Overall, our findings reveal that telomeres are protected in a privileged manner by FOXO3a during aging, supporting their role as central longevity regulators.

## Materials and Methods

### Cell culture

Human myoblasts in this study have been previously described (Robin *et al*, 2020). In brief, cells were platted in dishes coated with 0.1% pigskin gelatin in 4:1 Dulbecco modified Eagle medium/Medium 199 supplemented with 15% FBS, 0.02M HEPES, 1.4mg/l vitamin B12, 0.03mg/l ZnSO_4_, 0.055mg/l dexamethasone, 2.5μg/l hepatocyte growth factor and 10μg/l basic fibroblast growth factor. Cultures were passaged at ~70% confluency. Differentiation into myotubes was started following a change to differentiation medium (2% horse serum in 4:1 Dulbecco modified Eagle medium: Medium 199) when 90% confluent.

For infection of myotubes, myoblasts were seeded in 10 cm dishes, switch to differentiation media (2% Horse Serum) upon confluence (90%) and transduced at least week after. Cells were transduced at a MOI 2 using the different shRNAs.

BJ-HELT cells were immortalized by transduction of hTERT and SV40 ER genes (Biroccio *et al*, 2013). BjHeLT cells were grown in DMEM supplemented with 10% fetal bovine serum, penicillin (100 IU/ml), and streptomycin (100 μg/ml). Cells were cultured at 37°C, 5% CO2.

### Lentivirus infection and siRNA transfection

Lentiviruses were produced by transient calcium phosphate transfection of HEK293T cells with the virus packaging plasmids, p8.91 and pVSVg and a pWPIR-GFP lentiviral expression vector or pLKO plasmid, that contained the sequence of interest. We used MISSION shRNAs (pLKO.1; Sigma): *TERF2* (TRNC0000004812), *FOXO3A* (TRNC0000010335, only clone validated by Sigma MISSION shRNAs, to date) and control (SHC002). For upregulation, the pWPIR-GFP backbone was used and modified by Genscript to reach full-length FOXO3a pWPIR-FOXO3aWT-GFP and mutant forms pWPIR-ΔCR1-GFP, pWPIR-ΔFH-GFP, pWPIR-ΔCR2C-GFP, pWPIR-ΔCR3-GFP. pWPIR-TRF2-GFP and pWPIR-TRF2ΔB-GFP vectors have already been validated (Benarroch-Popivker *et al*, 2016). Transduction efficiency was determined by one-week puromycin treatment and count the number of resistant clones for pLKO.1 vectors and GFP-flow cytometry detection after 3 days for pWPIR-GFP vectors. siRNA transfections were performed with On-Target Plus SMARTpool (Dharmacon) and Dharmafect1 transfection reagent (T-2001, Dharmacon) for 72 hr.

Efficiency of each shRNA and siRNA was checked routinely by RT–qPCR or Western blotting.

### Cellular stressors treatments

BJ-HELT cells were treated with 250 μM H2O2 (Merk: 95294) for 24h. Bleomycin (Merk B2434), Paraquat (Merk: 36541) and Sodium Arsenite (Merk: S7400) were resuspended in distilled water and added to cell media at 50 μg/mL, 1,5 mM and 1 μM, respectively for 24h. For UV irradiation, media was removed, cells were washed with PBS and kept in a minimal volume of PBS. Then, cells were irradiated with UVA and UVB at 300 mJ/cm^2^ and let to recover for 2 hr before collection or fixation. To induce starvation, BJ-HELT cells were grown in normal conditions up to 70% confluency. Then, FBS containing media was removed, cells were washed with DMEM once and add DMEM without FBS. Cells were incubated under starvation for 1 week.

### Human biopsies and Ethic statement

The collection of fetal muscle biopsies was approved by the “Agence Française de la Biomedecine” of the Ministery of Health for legal access to the biological material in full accordance with the law (research protocol number PFS13-006). Samples were obtained after therapeutic abortion. Parents have provided written informed consent for the use of biopsies for medical research in accordance with the Declaration of Helsinki. Muscle biopsies were processed by fetopathologists from fetuses not affected by a muscular pathology. Skeletal muscle biopsies from teens and adults were obtained from the Nice Hospital (CHU l’Archet registered as protocol number DC-2015 2374) and from the Tumorothèque, Assistance Publique des Hôpitaux de Marseille, agreement n°AC-2013-1786, from healthy donors using standardized muscle biopsy protocol.

### Immunofluorescence-FISH and Telomere damage Induced Foci (TIFs)

Cells were grown onto glass coverslips or multichamber slides and fixed for 20 min with 3.7% formaldehyde. Cells were then permeabilized with 0.5% Triton X-100 for 15 min and dehydrated in increasing concentrations of ethanol for 3 min (50%, 70% and 100%). Hybridization of PNA probes was performed for at least 2 hr at RT after denaturation (5 min) in 70% formamide, 10 mM Tris pH 7.2 and 1% blocking solution (Roche) at 80°C. After that, the cells were washed in a 70% formamide, 10 mM Tris pH 7.2 solution for 30 min, followed by washes with 150 mM NaCl and 50 mM Tris pH 7.5 for 15 min. Next, the cells were incubated 1h with blocking buffer (3% BSA and 0,3% Triton X-100) and incubated overnight at 4°C with the desired antibody: FOXO3a, rabbit monoclonal (clone 75D8, 1:200, Cell Signaling Technology), 53BP1, rabbit polyclonal (NB 100-305, 1:200, Novus Biologicals) and 8-Oxoguanine, mouse monoclonal (clone 483.15, 1:50, MAB3560, Millipore). Cells were then washed with 1X PBS and incubated for 1 hr with the corresponding secondary antibody. Finally, slides were prepared with a DAPI containing mounting solution (Vectashield, Vector Laboratories).

### Immunofluorescence

Cells were grown onto glass coverslips or multichamber slides and fixed for 20 min with 3.7% formaldehyde. Cells were then permeabilized with 0.5% Triton X-100 for 15 min and blocked for 1h with (3% BSA and 0,3% Triton X-100). Next, cells incubated overnight at 4°C with the desired antibody: FOXO3a, rabbit monoclonal (clone 75D8, 1:200, Cell Signaling Technology). Cells were then washed with 1X PBS and incubated for 1 hr with the corresponding secondary antibody. Finally, slides were prepared with a DAPI containing mounting solution (Vectashield, Vector Laboratories).

### Reactive Oxygen Species (ROS)

Cells were grown on cover slides and treated as indicated in the manufacturer’s ROS kit instructions (Enzo-51011) and previously described. Transduced differentiated cells were washed twice and 1ml of fresh differentiation media was added, with or without EGCG (10mM) or H_2_O_2_ (100μM) and incubated for 30min at 37°C. After additional wash and media renewal (1ml), cells were incubated for 1 hour at 37°C with a solution composed 2X ROS detection and Oxidative stress reagent (5mM; dilution 1:2500). Cells were then washed with 1X PBS and directly mounted using 15μl of vectashield+DAPI, without any fixation. Pictures were taken using a DeltaVision Elite system (GE). An average of 100 stacks and 50 nuclei were taken per conditions. Images were then treated using IMARIS. Intensities of ROS foci and DAPI staining were used for analyses, excluding single-nuclei cells for myotubes analysis.

### Proximity Ligation Assay (PLA)

Cells were, fixed with 3,7% formaldehyde, permeabilized with 0.5% Triton X-100 for 15 min, and blocked with Duolink kit Blocking solution (DUO94001, Sigma-Aldrich). Cells were then incubated with primary antibodies to TRF2, mouse monoclonal, (NB100-56506, 1:300, Novus Biologicals), FOXO3a, rabbit monoclonal (clone 75D8, 1:1500, Cell Signaling Technology). Then, cells were incubated for 1h at 37°C with DNA-linked secondary antibodies (PLA probes), analyzed with the Duolink *In Situ* Red Mouse/Rabbit kit assay according to the manufacturer’s instructions (DUO94001, Sigma-Aldrich). In brief, the ligation and amplifications steps were performed for 30 min and 2 hr, respectively, at 37°C. Finally, cells were fixed again with formaldehyde 37°C. Images were taken with Deltavision Elite^®^ and analyzed with FIJI software. Only nuclear spots were considered for analysis. Each antibody used was first tested by immunofluorescence. PLA negative controls were performed to test each couple of antibodies: each antibody alone or without any antibody but with mouse/rabbit PLA probes.

### Chromatin-Immunoprecipitation (ChIP)

Samples were crosslinked for 10 min at RT and 20 min at 4°C with 0.8% formaldehyde (methanol free, ultrapure EM grade, Polysciences, Inc; Warrington PA). Reaction was stop at RT for 10 min with the addition of Glycine to a final concentration of 0.125 M. Cells were rinsed twice with ice-cold 1X PBS, scraped from the dish and pelleted after centrifugation (800g, 5min at 4°C). For sonication, we used a total processing time of 15min per sample in a Bioruptor (Diagenode) using the following settings: 14 cycles; 30 Sec ON/30 Sec OFF on High power. Sonicated DNA was controlled on a 2% agarose gel, adequate sonication is achieved when a smear ranging from 200-700bp is obtained. IPs were processed using a 4°C O/N incubation with TRF2 antibody at 1.5μg; Novus: NB100-56506); 1μl of each preparation: IP, IgG, Rabbit non-immune Serum. Next, magnetic beads (Dynabeads, Life Technologies) were added for 3 h. Samples were washed with a low salt buffer (150 mM NaCl, 1% Triton X-100, 0.1% SDS), a high salt buffer (500 mM NaCl, 1% Triton X-100, 0.1% SDS) and a lithium salt buffer (0.25 M LiCl, 1% NP40, 1% deoxycholic acid). Chromatin was eluted (1% SDS, 0.1 M NaHCO3 solution), and the cross-linked chromatin was reversed at 65°C overnight. The DNA was treated with RNaseA and proteinase K, followed by phenol–chloroform purification. The DNA obtained from ChIP was denatured and blotted onto nylon membranes using a slot blot apparatus, cross-linked, and hybridized with telomere and Alu repeats radioactively labeled probes. The membranes were exposed onto phosphorimager screens, and the signal intensity was quantified with ImageQuant software.

### Co-Immunoprecipitation

Skeletal muscle human biopsies were lysed in 50 mM Tris-HCl, 150 mM NaCl and 0,1% NP40; complemented with phosphatase inhibitors (Roche). Samples were sonicated using a Bioruptor (Diagenode) for 5 minutes (30 sec ON/ 30 sec OFF on High Power). Then, tubes were centrifugated 5 min at 4°C, 12 000 g. 10% of each sample was used for Input. Immunoprecipitations were processed using a TRF2 antibody (Novus: NB100-56506) at 4°C, O/N incubation. Next, magnetic beads (Dynabeads Protein G, Thermofisher, 10004D) were added for 1h at 4°C. Samples were washed with 50 mM Tris-HCl, 150 mM NaCl and 0,1% NP40; complemented with phosphatase inhibitors (Roche), five times. Resuspend samples in 2X Laemmli. Finally, samples were denaturated at 95°C for min and load then on a SDS-PAGE gel for Western Blot running.

### Western Blot

Cells were collected in 1X PBS and spin down (250g, 5min) and pellet stored at −80°C for further use. Whole cell lysates were prepared from cells by adding cell lysis RIPA buffer, complemented with phosphatase inhibitors (Roche) and kept 30 min on ice. For each sample, 30μg of protein was resolved in a 4-15% gradient mini-protean precast polyacrylamide gels (BioRad) and transferred to nitrocellulose membranes (Whatman, GE Healthcare) for 1h at 4°C. After blocking for 1 hour with 5% skim milk in PBST (0.1% Tween-20 in PBS), the membranes were incubated overnight at 4°C with primary antibodies diluted in 5% milk. The following primary antibodies were used: TRF2, mouse monoclonal, (NB100-56506, 1:1000, Novus Biologicals); TRF2, rabbit monoclonal, (NB110-57130, 1:5000, Novus Biologicals); FOXO3a, rabbit monoclonal (clone 75D8, 1:1500, Cell Signaling Technology); GAPDH, rabbit polyclonal (1:2000, NB100-56875, Novus Biologicals). The membranes were then rinsed three times in PBST for 10min and incubated 1 hour at room temperature with appropriate secondary antibodies diluted (1:5000) in 5% PBST-milk (e.g. anti-mouse HRP IgG and anti-rabbit HRP IgG, Vector Labotratories). Membranes were developed using the Luminata Forte HRP substrate (Millipore) and exposed in the Fusion Solo apparatus (Vilbert Lourmat).

### RT-qPCR

Myotubes RNA was extracted by Tri Reagent (Trizol; Sigma T9424), following manufacturer’s instructions. BJ-HELT and myoblasts were lysed (RNeasy plus kit (Qiagen: 74034)). Total RNA purified according to the manufacturer’s instructions and was quantified on a Nanodrop 1000 spectrophotometer (Thermo Scientific). For Reverse Transcription (RT) 500ng RNA was reverse transcribed High-Capacity RNA-to-cDNA Kit (Thermo Scientific). Each qPCR contained 5X diluted cDNA, 0.2 μM primers, and SYBR Green Master Mix (Roche, 4913914 001). Quantitative RT-PCR (qRT-PCR) was permorfed in triplicates using FastStart universal SYBR Green master Mix (Roche). Melting curves were analyzed (SYBR green) to exclude nonspecific amplification products. We confirmed amplicon size at least once on agarose gels. Crossing-threshold (Ct) values were normalized by subtracting the geometric mean of two housekeeping genes (GAPDH and HPRT).

### Statistical Analysis

Statistical analysis was performed using the Prism 7 software (GraphPad). Quantitative data are displayed as means ± standard error of the mean. For comparison of two groups, we used two-tailed Mann–Whitney U-test or Student’s t-test, and for multiple groups, the Kruskal–Wallis test was used. *p-value* < 0.05 were considered significant (\**p* < 0.05, \*\**p* < 0.001, \*\*\**p* < 0.001, \*\*\*\**p* < 0.0001).

## Acknowledgments and funding

The IRCAN’s Molecular and Cellular Core Imaging (PICMI) is supported by “le Cancéropole PACA, la Région Provence Alpes-Côte d’Azur, le Conseil Départemental 06”, and INSERM.

This work was supported by the cross-cutting INSERM program on aging (AGEMED), ‘‘Investments for the Future’’ LABEXSIGNALIFE (reference ANR-11-LABX-0028-01), the “Fondation ARC pour la recherche contre le cancer”, INCa (project REPLITOP), the ANR (projects TELOPOST and TELOCHROM) and the FRM (FDT202012010648).

## Author contributions

The project was design by EG, MSJB and JDR. Investigation and validation were performed by MSJB and JDR. SB, MM, ED and LM provided technical help. FM and SS provided human skeletal muscle biopsies. MSJB and EG wrote and edit the original draft. EG supervised this work. All authors read and commented on the manuscript.

## Data and materials availability

All data are available in the main text or the supplementary materials.

## Expanded view legends

**EV1.**
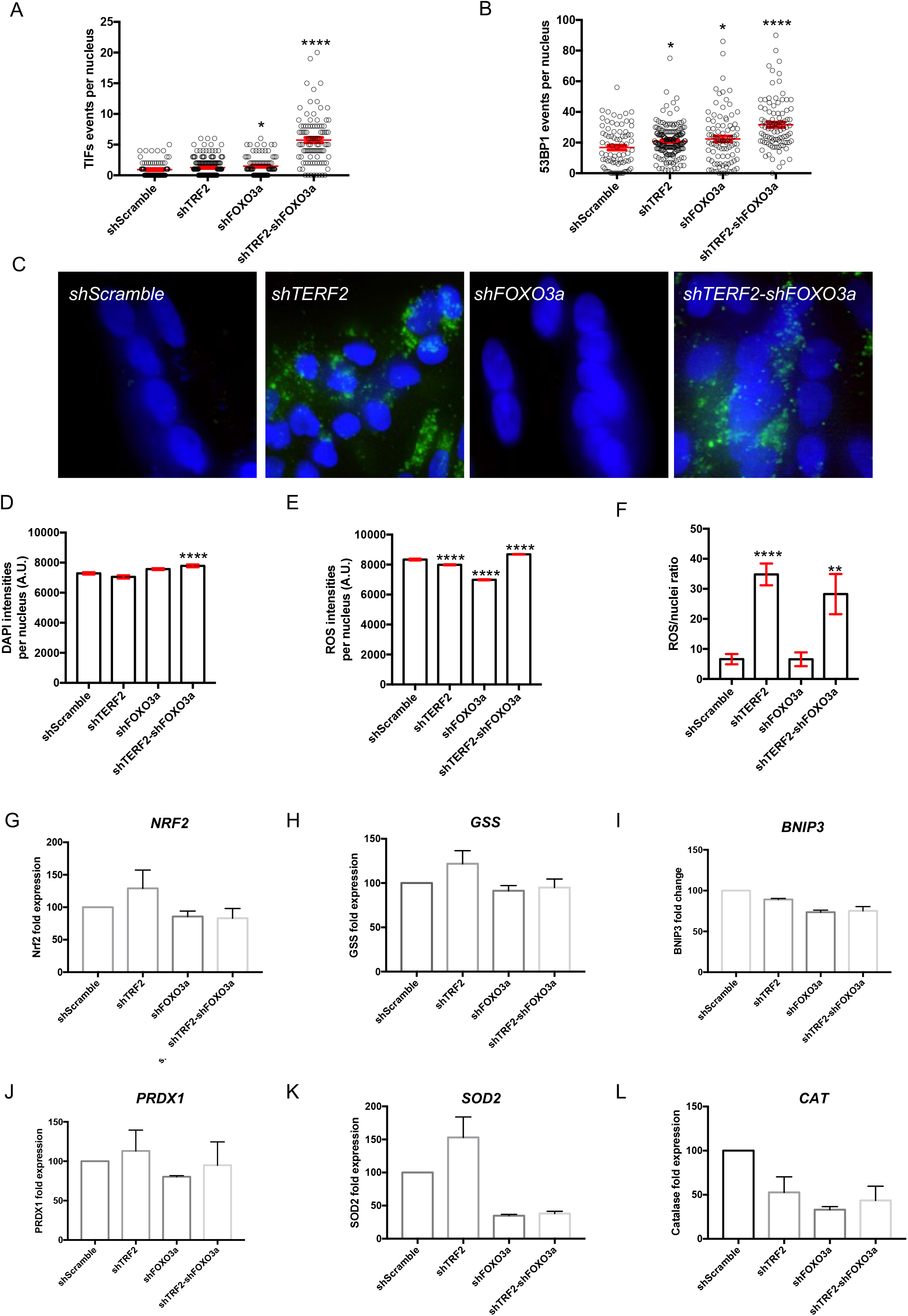
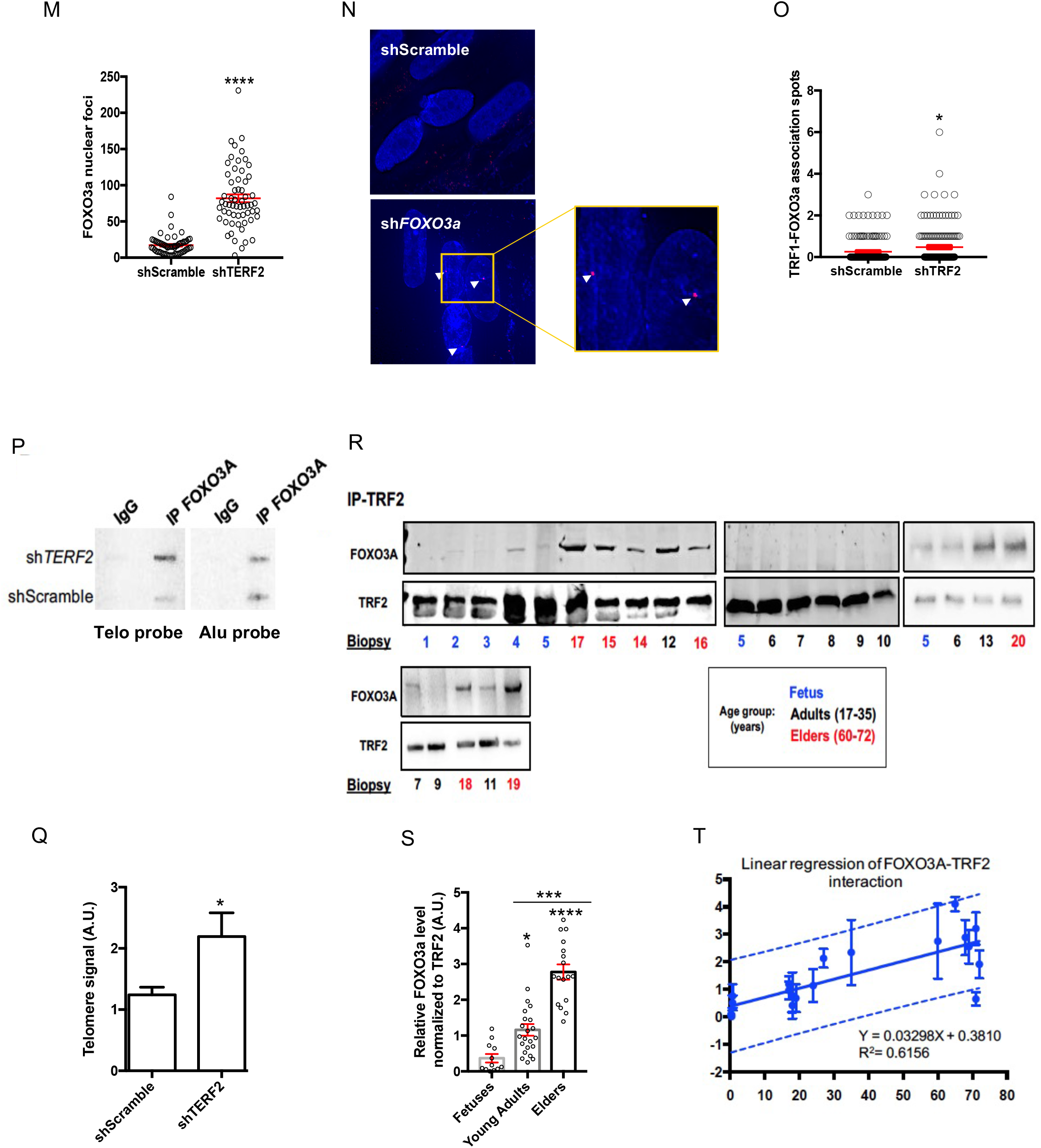
**a) Telomere damage is not due to FOXO3a inhibition induced oxidative stress.** (A) Telomere damage Induced Foci (TIF) and 53BP1. (B) quantification. Means ± SEM are shown. Only foci within multinucleated cells, that is, corresponding to postmitotic myotubes were counted. 30-40 nuclei were analyzed per replicate and per condition, n=4. (C) Representative images of ROS foci in transduced myotubes. (D) DAPI and ROS (E) intensities quantification in transduced myotubes represented in (C). On average, n>300 nuclei per conditions, means ± SEM are shown. (F) Total number of ROS foci normalized to the number of nuclei. Down-regulation of TERF2 significantly increases ROS. Statistical analyses were performed using oneway ANOVA test (\**p* < 0.05, \*\**p* < 0.001, \*\*\**p* < 0.001, \*\*\*\**p* < 0.0001). (G) to L) Gene expression quantified by RT-qPCR in transduced myotubes. Each measure represents the average fold-change expression of six independent repetitions (Biological triplicate in technical RT duplicate) normalized to two housekeeping genes (HKG: HPRT and GAPDH); Pfaffl method). Means ± SEM are shown. **b) FOXO3a directly binds to telomeres in myotubes, upon TRF2 downregulation**. (M) FOXO3a nuclear Foci quantification in transduced myotubes. *p-value* < 0.0001, Mann-Whitney test. (N) PLA TRF1-FOXO3a (red spots) in myotubes downregulated for TRF2. White arrows point association spots. Zoom at the left. (O) Quantification of two biological replicates. *p-value=* 0,0295. (P) and (Q) Slot blot of ChIP samples using a Telomeric (right) and Alu (left) probe in transduced myotubes and associated quantification normalized to Alu repeats. IgG shown as control. n=3 per condition; means ± SEM are shown. FOXO3A localizes to telomeres upon *TERF2* down-regulation (shScramble vs. *shTERF2*, p=0.0286; Man-Whitney test, α=0.05). (R) to (T) Immunoprecipitation (IP) using TRF2 antibody in human skeletal muscle biopsies, using TRF2 antibody for precipitation and FOXO3A for revelation. The TRF2-FOXO3A interaction enhances with ageing. Correlation and goodness of fit associated (R2), 95% interval confidence (blue lines) and means ± SEM are shown.

**EV2.**
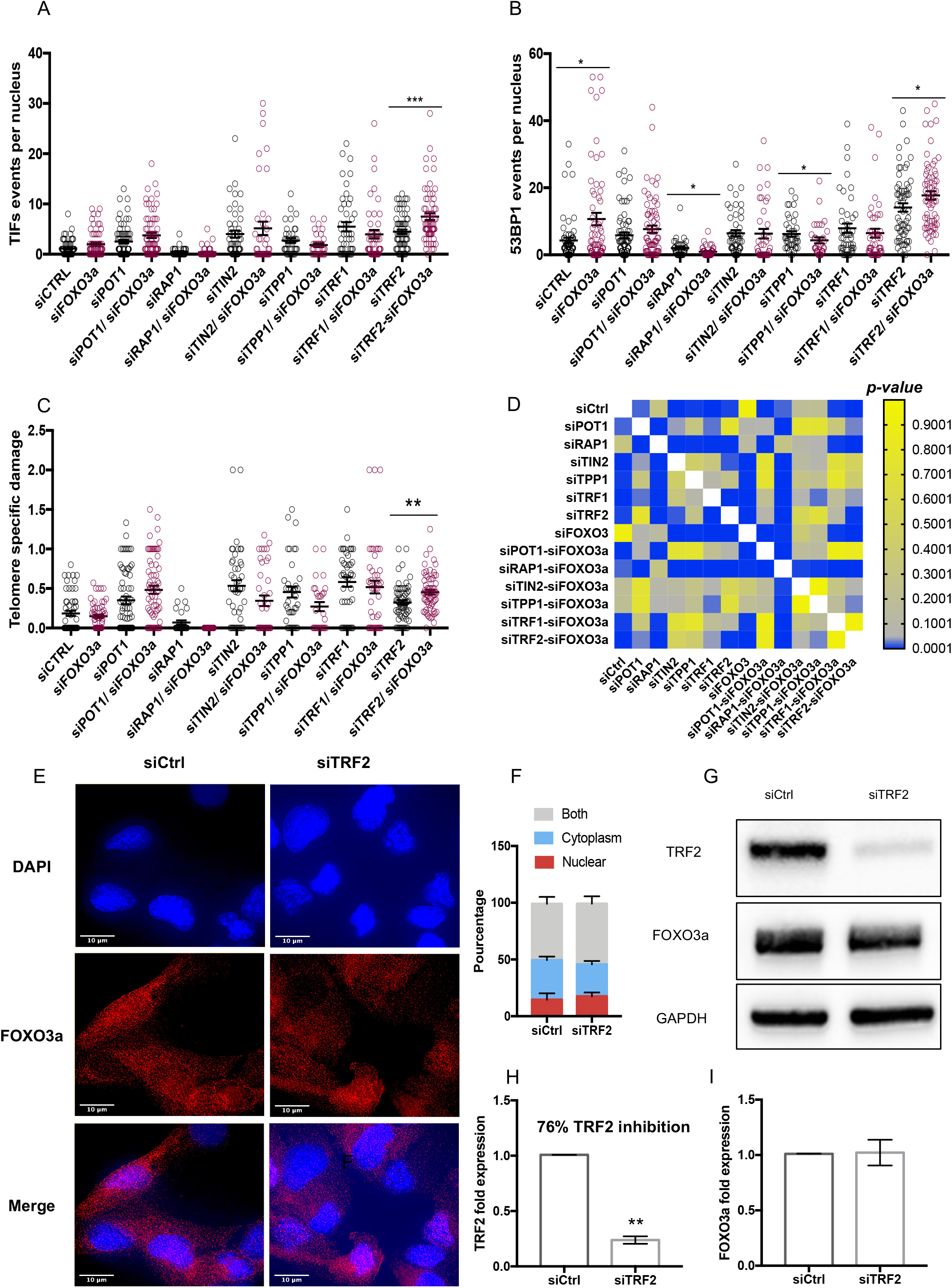
FOXO3a protects telomeres specifically upon *TERF2* inhibition in BJ-HELT cells. (A) TIFs in BJ-HELT fibroblasts transfected with siRNA against each shelterin protein (TRF1, TRF2, POT1, RAP1, TIN2 and TPP1) individually or combined to siFOXO3a. (B) 53BP1 foci per nucleus in conditions described in A). Statistical analysis was performed between single *vs* couple siRNA (example: siTRF1 *vs* siTRF1-siFOXO3a), using Mann–Whitney U-test (\**p* < 0.05, \*\**p* < 0.001, \*\*\**p* < 0.001). (C) Telomere specific damage of A and B (TIFs/ 53BP1). (D) Heat map of (C) statistical analysis using Mann–Whitney U-test. Conditions labelled in blue are significantly different, the threshold α= 0.05 *p-value* is represented in grey, nonsignificant conditions correspond to the yellow staining. (E) and (F) FOXO3a immunofluorescence and qualitatively determination of FOXO3a cellular localization: mostly nuclear, mostly cytoplasmic, nuclear and cytoplasmic. Approximatively 60 cells were analyzed per condition in biological triplicates. (G) Western Blotting showing FOXO3a protein levels in BJ-HELT cells downregulated forTRF2. (H) Downregulation of *TERF2* was also assessed by RT-qPCR. *p-values* of n=5 were obtained using the Mann-Whitney test (* p< 0.05). (I) FOXO3a expression measured by RT-qPCR upon TRF2 inhibition.

**EV3.**
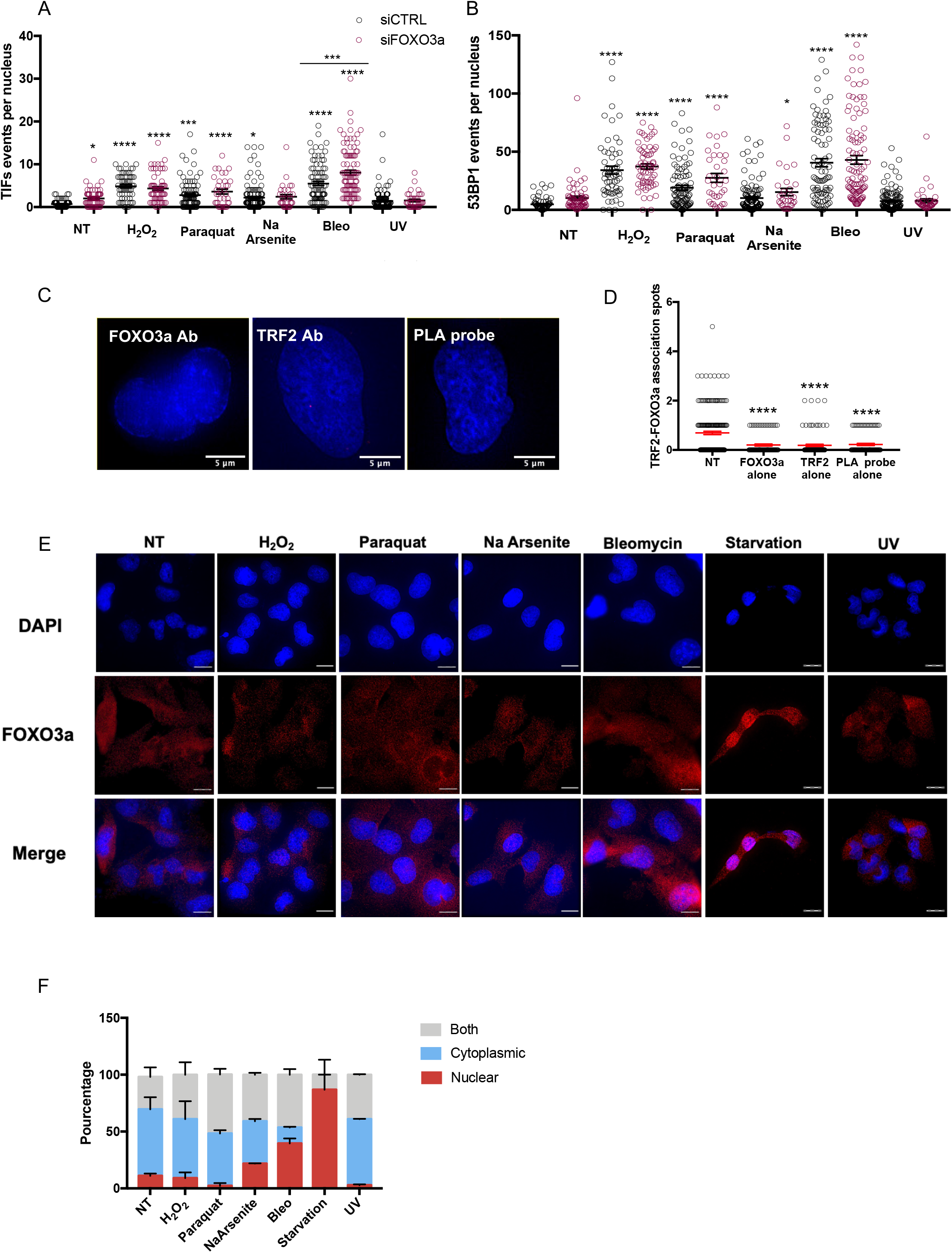
FOXO3a-mediated telomere protection upon cellular stress. (A) TIFs and 53BP1 (B) in transfected BJ-HELT cells with siControl (black circles) or siFOXO3a (pink circles) and exposed to different stressors. Statistical analyses were performed using Kruskal-Wallis’ multiple comparisons test (\**p* < 0.05, \*\**p* < 0.001, \*\*\**p* < 0.001, \*\*\*\**p* < 0.0001) in biological triplicates. (C) Representative images of PLA negative controls: FOXO3a antibody alone + PLA probe, TRF2 antibody alone + PLA probe, no antibodies but PLA probe alone. (D) PLA negative control quantification, compared to PLA (TRF2, FOXO3a antibodies and PLA probes added). Kruskal-Wallis’ multiple comparisons test (\*\*\*\**p* < 0.0001) in three biological replicates. (E) FOXO3a cellular localization upon different kind of stress. FOXO3a pattern was qualitatively determined as mostly nuclear, mostly cytoplasmic, nuclear and cytoplasmic in (F). Approximatively 30 cells were analyzed per condition in biological duplicates.

**EV4.**
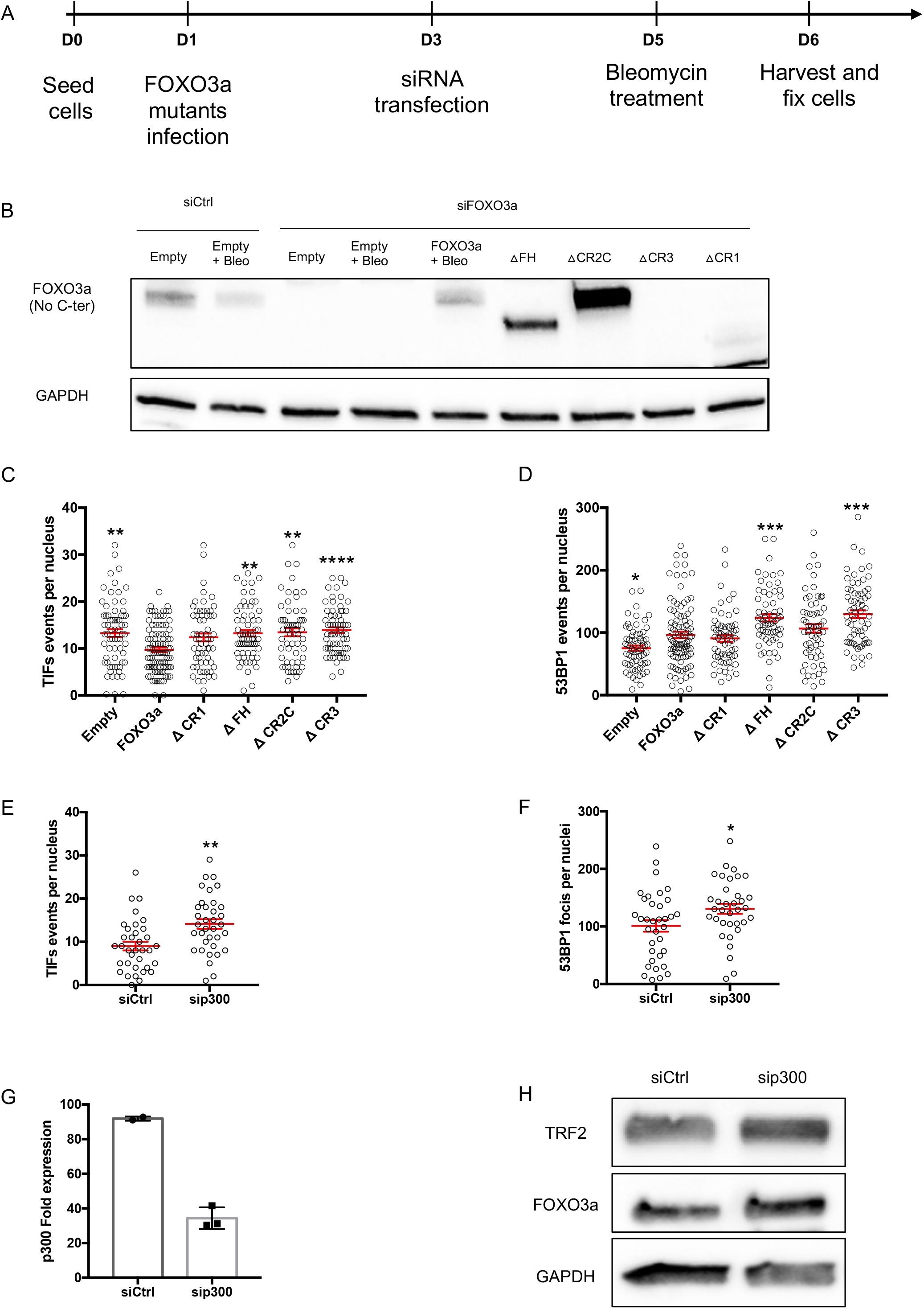
p300 is involved in the FOXO3a-dependent telomere protection. (A) FOXO3a truncated mutant overexpression and endogen FOXO3a downregulation technical strategy. Cells were platted at day 0 and transduced at MOI 2 with lentiviral FOXO3a full-length or truncated forms and Empty vector at day 1. At day 3, each condition was transfected with siControl or siFOXO3a. At day 5, bleomycin (50 μg/mL) was added and cells were fixed or harvest 24hr later. (B) Western blotting showing endogen FOXO3a downregulation, FOXO3a full-length rescue and expression of FOXO3a truncated proteins. FOXO3a-Abcam ab12162 polyclonal antibody recognizes 653-668 aa of human FOXO3a (C-terminal), does not recognizes ΔCR3. Cells were transduced 5 days with lentiviruses and the last 3 days transfected with siRNA. (C) TIFs and (D) 53BP1 in BJ-HELT cells transfected with siFOXO3a and transduced with the truncated FOXO3a overexpression. All conditions were treated with bleomycin (50 μg/mL, 24hr), n=3. Conditions were compared to siFOXO3a-FOXO3a+Bleo using a Kruskal-Wallis’ multiple comparisons test (\**p* < 0.05, \*\**p* < 0.001, \*\*\**p* < 0.001, \*\*\*\**p* < 0.0001). (E) TIFs and (F) 53BP1 in BJ-HELT cells transfected with sip300 and treated with bleomycin. Man-Whitney U test (*p-value=* 0,0011 and 0,0368, respectively). (G) p300 expression measured by RT-qPCR to control p300 inhibition by siRNA (n=3). (H) Western blotting showing that neither TRF2 nor FOXO3a expression are affected by p300 KD.

## References

Abdallah P, Luciano P, Runge KW, Lisby M, Géli V, Gilson E & Teixeira MT (2009) A two-step model for senescence triggered by a single critically short telomere. Nat Cell Biol 11: 988–993

Adamowicz M, Vermezovic J & d’Adda di Fagagna F (2016) NOTCH1 Inhibits Activation of ATM by Impairing the Formation of an ATM-FOXO3a-KAT5/Tip60 Complex. Cell Rep 16: 2068–2076

Alder JK, Barkauskas CE, Limjunyawong N, Stanley SE, Kembou F, Tuder RM, Hogan BLM, Mitzner W & Armanios M (2015) Telomere dysfunction causes alveolar stem cell failure. PNAS 112: 5099–5104

Armanios M & Blackburn EH (2012) The telomere syndromes. Nat Rev Genet 13: 693–704

Augereau A, Mariotti M, Pousse M, Filipponi D, Libert F, Beck B, Gorbunova V, Gilson E & Gladyshev VN (2021) Naked mole rat TRF1 safeguards glycolytic capacity and telomere replication under low oxygen. Sci Adv 7: eabe0174

Barnes RP, Fouquerel E & Opresko PL (2019) The impact of oxidative DNA damage and stress on telomere homeostasis. Mech Ageing Dev 177: 37–45

Benarroch-Popivker D, Pisano S, Mendez-Bermudez A, Lototska L, Kaur P, Bauwens S, Djerbi N, Latrick CM, Fraisier V, Pei B, et al (2016) TRF2-Mediated Control of Telomere DNA Topology as a Mechanism for Chromosome-End Protection. Mol Cell 61: 274–286

Biroccio A, Cherfils-Vicini J, Augereau A, Pinte S, Bauwens S, Ye J, Simonet T, Horard B, Jamet K, Cervera L, et al (2013) TRF2 inhibits a cell-extrinsic pathway through which natural killer cells eliminate cancer cells. Nat Cell Biol 15: 818–828

Brunet A, Sweeney LB, Sturgill JF, Chua KF, Greer PL, Lin Y, Tran H, Ross SE, Mostoslavsky R, Cohen HY, et al (2004) Stress-dependent regulation of FOXO transcription factors by the SIRT1 deacetylase. Science 303: 2011–2015

Cherfils-Vicini J, Iltis C, Cervera L, Pisano S, Croce O, Sadouni N, Gyõrffy B, Collet R, Renault VM, Rey-Millet M, et al (2019) Cancer cells induce immune escape via glycocalyx changes controlled by the telomeric protein TRF2. EMBO J 38

Cubiles MD, Barroso S, Vaquero-Sedas MI, Enguix A, Aguilera A & Vega-Palas MA (2018) Epigenetic features of human telomeres. Nucleic Acids Res 46: 2347–2355

De Bonis ML, Ortega S & Blasco MA (2014) SIRT1 Is Necessary for Proficient Telomere Elongation and Genomic Stability of Induced Pluripotent Stem Cells. Stem Cell Reports 2: 690–706

Epel ES, Blackburn EH, Lin J, Dhabhar FS, Adler NE, Morrow JD & Cawthon RM (2004) Accelerated telomere shortening in response to life stress. Proc Natl Acad Sci U S A 101: 17312–17315

Foley NM, Hughes GM, Huang Z, Clarke M, Jebb D, Whelan CV, Petit EJ, Touzalin F, Farcy O, Jones G, et al (2018) Growing old, yet staying young: The role of telomeres in bats’ exceptional longevity. Science Advances 4: eaao0926

Gilson E & Géli V (2007) How telomeres are replicated. Nat Rev Mol Cell Biol 8: 825–838

Gorgoulis V, Adams PD, Alimonti A, Bennett DC, Bischof O, Bishop C, Campisi J, Collado M, Evangelou K, Ferbeyre G, et al (2019) Cellular Senescence: Defining a Path Forward. Cell 179: 813–827

Her YR & Chung IK (2013) p300-mediated acetylation of TRF2 is required for maintaining functional telomeres. Nucleic Acids Res 41: 2267–2283

Jacome Burbano MS & Gilson E (2021) The Power of Stress: The Telo-Hormesis Hypothesis. Cells 10: 1156

Jones OR, Scheuerlein A, Salguero-Gómez R, Camarda CG, Schaible R, Casper BB, Dahlgren JP, Ehrlén J, García MB, Menges ES, et al (2014) Diversity of ageing across the tree of life. Nature 505: 169–173

Kim W, Ludlow AT, Min J, Robin JD, Stadler G, Mender I, Lai T-P, Zhang N, Wright WE & Shay JW (2016) Regulation of the Human Telomerase Gene TERT by Telomere Position Effect—Over Long Distances (TPE-OLD): Implications for Aging and Cancer. PLOS Biology 14: e2000016

Kishi S, Bayliss PE, Uchiyama J, Koshimizu E, Qi J, Nanjappa P, Imamura S, Islam A, Neuberg D, Amsterdam A, et al (2008) The identification of zebrafish mutants showing alterations in senescence-associated biomarkers. PLoS Genet 4: e1000152

de Lange T (2018) Shelterin-Mediated Telomere Protection. Annu Rev Genet 52: 223–247

López-Otín C, Blasco MA, Partridge L, Serrano M & Kroemer G (2013) The Hallmarks of Aging. Cell 153: 1194–1217

Mammucari C, Milan G, Romanello V, Masiero E, Rudolf R, Del Piccolo P, Burden SJ, Di Lisi R, Sandri C, Zhao J, et al (2007) FoxO3 controls autophagy in skeletal muscle in vivo. Cell Metab 6: 458–471

Martínez G, Duran-Aniotz C, Cabral-Miranda F, Vivar JP & Hetz C (2017) Endoplasmic reticulum proteostasis impairment in aging. Aging Cell 16: 615–623

Martins R, Lithgow GJ & Link W (2016) Long live FOXO: unraveling the role of FOXO proteins in aging and longevity. Aging Cell 15: 196–207

Milan G, Romanello V, Pescatore F, Armani A, Paik J-H, Frasson L, Seydel A, Zhao J, Abraham R, Goldberg AL, et al (2015) Regulation of autophagy and the ubiquitin–proteasome system by the FoxO transcriptional network during muscle atrophy. Nat Commun 6: 6670

Morgan RG, Walker AE, Trott DW, Machin DR, Henson GD, Reihl KD, Cawthon RM, Denchi EL, Liu Y, Bloom SI, et al (2019) Induced Trf2 deletion leads to aging vascular phenotype in mice associated with arterial telomere uncapping, senescence signaling, and oxidative stress. J Mol Cell Cardiol 127: 74–82

Muñoz-Lorente MA, Cano-Martin AC & Blasco MA (2019) Mice with hyper-long telomeres show less metabolic aging and longer lifespans. Nat Commun 10: 1–14

Nakae J, Cao Y, Daitoku H, Fukamizu A, Ogawa W, Yano Y & Hayashi Y (2006) The LXXLL motif of murine forkhead transcription factor FoxO1 mediates Sirt1-dependent transcriptional activity. J Clin Invest 116: 2473–2483

Palacios JA, Herranz D, De Bonis ML, Velasco S, Serrano M & Blasco MA (2010) SIRT1 contributes to telomere maintenance and augments global homologous recombination. J Cell Biol 191: 1299–1313

Partridge L & Barton NH (1993) Optimally, mutation and the evolution of ageing. Nature 362: 305–311

Robin JD, Jacome Burbano M-S, Peng H, Croce O, Thomas JL, Laberthonniere C, Renault V, Lototska L, Pousse M, Tessier F, et al (2020) Mitochondrial function in skeletal myofibers is controlled by a TRF2-SIRT3 axis over lifetime. Aging Cell 19: e13097

Sahin E, Colla S, Liesa M, Moslehi J, Müller FL, Guo M, Cooper M, Kotton D, Fabian AJ, Walkey C, et al (2011) Telomere dysfunction induces metabolic and mitochondrial compromise. Nature 470: 359–365

Sandri M, Sandri C, Gilbert A, Skurk C, Calabria E, Picard A, Walsh K, Schiaffino S, Lecker SH & Goldberg AL (2004) Foxo Transcription Factors Induce the Atrophy-Related Ubiquitin Ligase Atrogin-1 and Cause Skeletal Muscle Atrophy. Cell 117: 399–412

Schumacher B, Pothof J, Vijg J & Hoeijmakers JHJ (2021) The central role of DNA damage in the ageing process. Nature 592: 695–703

Senf SM, Sandesara PB, Reed SA & Judge AR (2011) p300 Acetyltransferase activity differentially regulates the localization and activity of the FOXO homologues in skeletal muscle. Am J Physiol Cell Physiol 300: C1490–C1501

Sharma S, Mukherjee AK, Roy SS, Bagri S, Lier S, Verma M, Sengupta A, Kumar M, Nesse G, Pandey DP, et al (2021) Human telomerase is directly regulated by non-telomeric TRF2-G-quadruplex interaction. Cell Reports 35: 109154

Singh PP, Demmitt BA, Nath RD & Brunet A (2019) The Genetics of Aging: A Vertebrate Perspective. Cell 177: 200–220

Soerensen M, Nygaard M, Dato S, Stevnsner T, Bohr VA, Christensen K & Christiansen L (2015) Association study of FOXO3A SNPs and aging phenotypes in Danish oldest-old individuals. Aging Cell 14: 60–66

Tran H, Brunet A, Grenier JM, Datta SR, Fornace AJ, DiStefano PS, Chiang LW & Greenberg ME (2002) DNA repair pathway stimulated by the forkhead transcription factor FOXO3a through the Gadd45 protein. Science 296: 530–534

Tsai W-B, Chung YM, Takahashi Y, Xu Z & Hu MC-T (2008) Functional interaction between FOXO3a and ATM regulates DNA damage response. Nat Cell Biol 10: 460–467

Wagner K-D, Ying Y, Leong W, Jiang J, Hu X, Chen Y, Michiels J-F, Lu Y, Gilson E, Wagner N, et al (2017) The differential spatiotemporal expression pattern of shelterin genes throughout lifespan. Aging (Albany NY) 9: 1219–1228

Wang F, Marshall CB, Yamamoto K, Li G-Y, Gasmi-Seabrook GMC, Okada H, Mak TW & Ikura M (2012) Structures of KIX domain of CBP in complex with two FOXO3a transactivation domains reveal promiscuity and plasticity in coactivator recruitment. PNAS 109: 6078–6083

Wang F, Marshall CB, Yamamoto K, Li G-Y, Plevin MJ, You H, Mak TW & Ikura M (2008) Biochemical and Structural Characterization of an Intramolecular Interaction in FOXO3a and Its Binding with p53. Journal of Molecular Biology 384: 590–603

Webb AE & Brunet A (2014) FOXO transcription factors: key regulators of cellular quality control. Trends in Biochemical Sciences 39: 159–169

White RR, Maslov AY, Lee M, Wilner SE, Levy M & Vijg J (2020) FOXO3a acts to suppress DNA double-strand break-induced mutations. Aging Cell 19: e13184

Wirthlin M, Lima NCB, Guedes RLM, Soares AER, Almeida LGP, Cavaleiro NP, Loss de Morais G, Chaves AV, Howard JT, Teixeira M de M, et al (2018) Parrot Genomes and the Evolution of Heightened Longevity and Cognition. Curr Biol 28: 4001–4008.e7

Wright WE, Piatyszek MA, Rainey WE, Byrd W & Shay JW (1996) Telomerase activity in human germline and embryonic tissues and cells. Dev Genet 173–179

Ye J, Renault VM, Jamet K & Gilson E (2014) Transcriptional outcome of telomere signalling. Nature Reviews Genetics 15: 491

Zhao J, Brault JJ, Schild A, Cao P, Sandri M, Schiaffino S, Lecker SH & Goldberg AL (2007) FoxO3 coordinately activates protein degradation by the autophagic/lysosomal and proteasomal pathways in atrophying muscle cells. Cell Metab 6: 472–483

